# Application of nanopore sequencing for accurate identification of bacterial colonies

**DOI:** 10.1101/2023.01.03.522650

**Authors:** Austin Marshall, Daniel T. Fuller, Paul Dougall, Kavindra Kumaragama, Suresh Dhaniyala, Shantanu Sur

## Abstract

Culture based detection remains to be one of the most reliable and acceptable techniques to detect extremely low quantity pathogens present in a sample. The process typically involves inoculating the sample on an agar plate to allow growth of the microorganisms to form colonies, followed by the identification of the individual colonies, commonly by DNA sequencing of a PCR-amplified targeted gene. Sanger method is often the default choice of sequencing as it offers affordable and accurate results for a single species. However, the technique could pose limitations in certain situations such as identification of multi-species microbial colonies. In this work, we compared the performance of Sanger sequencing with MinION nanopore sequencing in identifying bacterial colonies derived from bioaerosol samples. We conducted Sanger and nanopore sequencing of full-length 16S rRNA genes from seven bacterial colonies derived from bioaerosol samples and compared the outcome by alignment against NCBI 16S reference database. We found that for five out of seven colonies both techniques indicated the presence of the same bacterial genus. For one of the remaining colonies, a noisy Sanger electropherogram failed to generate a meaningful sequence, but nanopore sequencing identified it to be a mix of two bacterial genera *Alkalihalobacillus* and *Kocuria*. For the other remaining colony, the Sanger sequencing suggested *Micrococcus* with a clean electropherogram, however, the nanopore sequencing suggested the presence of an additional genus *Paraburkholderia*. Further corroborating these findings with mock multispecies colonies from pure bacterial DNA samples, we confirm that nanopore sequencing is comparable to the Sanger method in identifying colonies with single bacterial species but is the superior method in classifying individual bacterial components with their relative abundances in multispecies colonies. Our results suggest that nanopore sequencing could be advantageous over Sanger sequencing for colony identification in culture-based analysis of environmental samples such as bioaerosol where direct inoculation of the sample to culture plate might lead to formation of multispecies colonies.

## Introduction

Culture-based methods are still routinely used and are considered a reliable technique for the detection of pathogens and other viable microorganisms in many scientific fields, despite the recent advancement in genomic and other culture-independent rapid detection methods.^1–5^ For example, in food safety monitoring, culturing is often used to identify and quantify potential food-borne pathogens.^6^ Clinical samples with suspected infections are routinely cultured to detect bacterial and fungal pathogens.^7,8^ Culture is also used commonly for microbial analysis of environmental samples such as soil, water, and more recently within air due to their extremely low concentrations.^2,9–11^

Correct identification of microbial colonies grown on agar plates is a key step in culture-based detection strategies. While it was done in the past using a battery of biochemical tests, they have since been mostly replaced by sequencing-based approaches. Targeted amplicon analysis is a commonly adapted approach for taxonomic classification and for studying phylogenetic relationships due to the conserved nature of many essential genes. The 16S ribosomal RNA gene (henceforth abbreviated as 16S for brevity), contains nine variable regions and is present universally in bacteria and archaea, providing a robust tool for the classification of bacteria and archaea.^12–16^ While 16S amplicon analysis is widely used for identification, the implementation of different sequencing technologies can influence the resolution of the results and the scope of application.

Sanger sequencing, first introduced more than four decades ago, is still widely used, and remains one of the primary sequencing tools for the identification of microbial colonies through targeted gene amplification. Sanger sequencing is highly accurate in sequencing reads (up to ∼1000 bases) and is often held as a reference or standard to compare the accuracy of other sequencing techniques.^17–20^ One major limitation of Sanger sequencing is that only one homogenous DNA sequence can be read by this technique, and the presence of additional sequences in the sample will impact the output.^12,13^ Thus Sanger sequencing is not suitable for sequencing 16S amplicons from a sample with mixed microbial composition. For such applications, next-generation sequencing (NGS), which employs massively parallel short-read sequencing, is commonly used.

Although microbes in nature exist in rich and diverse communities, microbes derived from such samples after culturing enables the possibility of studying individually. The colonies are usually homogeneous as they are generated by aggregated growth from a single cell, however, multispecies colonies are reported where distant bacterial species associate in a single colony structure with specific interactions observed between them.^21,22^ Multispecies colonies n potentially form when environmental samples, specifically bioaerosol samples are directly plated on agar media. Microbes in the air are reported to exist as aggregates of variable size, often bound to particulate matter.^23,24^ Since conducting culture experiments after capturing bioaerosol particles directly on an agar plate (e.g., by depositional sampling, impaction) is a commonly accepted practice, viable bacterial aggregates in the air can lead to the formation of multispecies colonies in these cultures.^5^ For such colonies, Sanger sequencing will be a less ideal method for amplicon-based identification as the technique will not be able to resolve the identity of individual bacteria. 16S analysis using NGS technique such as Illumina-based sequencing can be used in this situation to correctly classify all bacterial taxa from a multispecies colony; however, owing to the limitation of short reads, NGS allows only a segment (<500 bp) of the full 16S gene to be sequenced and this restricts the resolution of taxonomic classification up to genus level.^25–27^

Third-generation sequencing is commonly referred to the sequencing platforms offered by Oxford Nanopore Sequencing (ONT) and Pacific Biosciences (PacBio), which overcomes some of the major limitations of NGS by enabling long-read sequencing.^28,29^ Among these techniques, MinION nanopore sequencing from ONT utilizes a protein nanopore complex to guide a DNA strand to translocate through the pore and determines the sequence from the changes in ionic conductivities as different nucleotide bases pass through the pore.^29^ Nanopore sequencing made remarkable advancements in the last decade with significant improvement in sequencing accuracy and capacity, which combined with an extremely portable, inexpensive sequencing device, and relatively simple library preparation procedures have attracted interest in a wide range of sequencing applications.^30–34^ The long-read capability of nanopore sequencing allows for full-length 16S gene amplicon sequencing with the ability to discriminate up to species level in a sample of mixed bacterial composition.^27^ Furthermore, multiplexing the samples by barcoding enables running multiple samples on a single run, enhancing the throughput, and reducing the cost. Together, these features of nanopore sequencing make it a potentially attractive procedure for the identification of bacterial colonies through 16S amplicon analysis.

In this study, we compared Sanger and nanopore sequencing for the identification of bacterial colonies derived from bioaerosol samples and explored any advantages afforded by nanopore sequencing. Targeted amplification of full-length 16S genes was conducted for individual colonies, and the amplicons were sequenced using both techniques. We investigated the accuracy of these two techniques on colony identification, especially when there is a potential for the existence of multispecies colonies. The findings were further corroborated with mock multispecies colony samples.

## Results

### 16S sequencing and colony identification: Sanger method

In this study, we used seven distinct and randomly selected bacterial colonies obtained by culturing environmental bioaerosol samples on Tryptic Soy Agar (TSA). Genomic DNA was extracted from each of these colonies and Sanger sequencing was performed on full-length 16S PCR amplification products (Fig.1).

**Figure 1.**
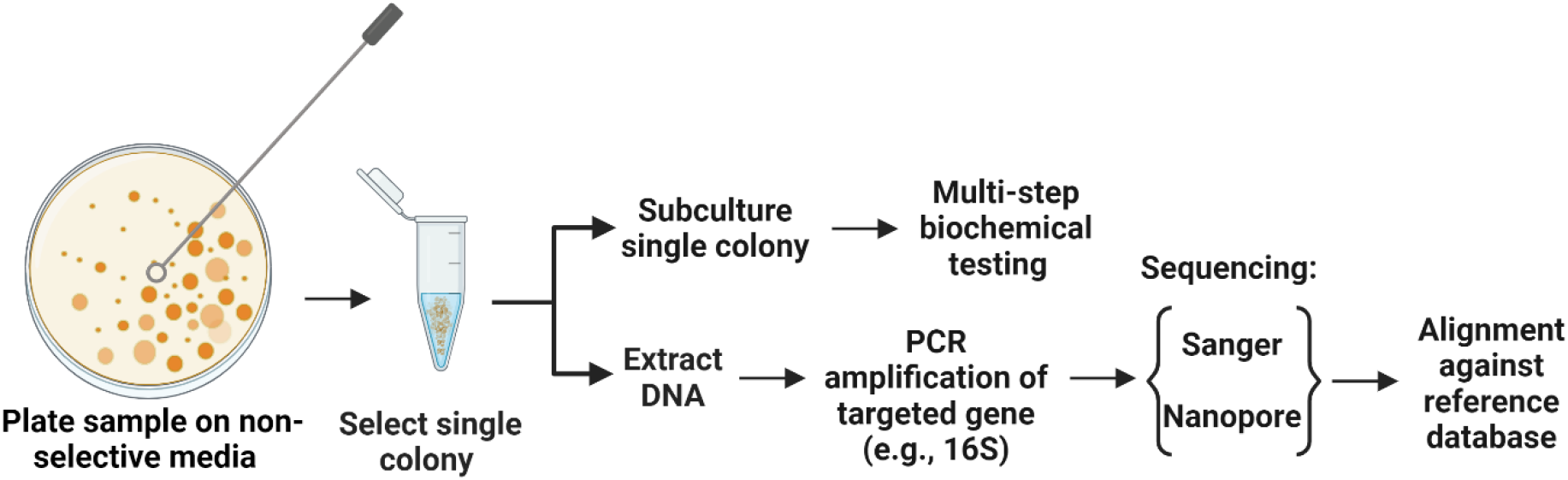
Schema showing two main approaches for the identification of bacterial colonies. Both Sanger and nanopore sequencing can be used for identification by targeted gene amplification.

The results of Sanger sequencing are summarized in Figure 2. The sequencing generated a single consensus sequence for each colony. The sequences were identified by aligning against the NCBI 16S database using the nucleotide Basic Local Alignment Search Tool (BLASTn). The top match from the search was used for taxonomic classification and a genus level identification was assigned when there was at least 95% identity.^25^ Using this criterion, six out of seven colonies (colony **2**-**7**, Fig. 2) were successfully identified at genus level, however, no classification was possible for colony **1**. To understand why the sequence from colony **1** failed to provide a taxonomic classification, we looked at the electropherograms. The electropherograms from Sanger sequencing were used to assess the quality of the consensus sequence generated. A peak in an electropherogram represents the signal from a nucleotide base and is used to determine the base at a specific location of the consensus sequence. An electropherogram with high quality will consist of single, discrete peaks, while areas of poorer quality contain multiple overlapping peaks. We found that the electropherogram for colony **1** is considerably noisier than the electropherograms for other colonies with many overlapping peaks distributed throughout. The noisy electropherogram of colony **1** explains why a consensus sequence could not be obtained from Sanger sequencing.

**Figure 2.**
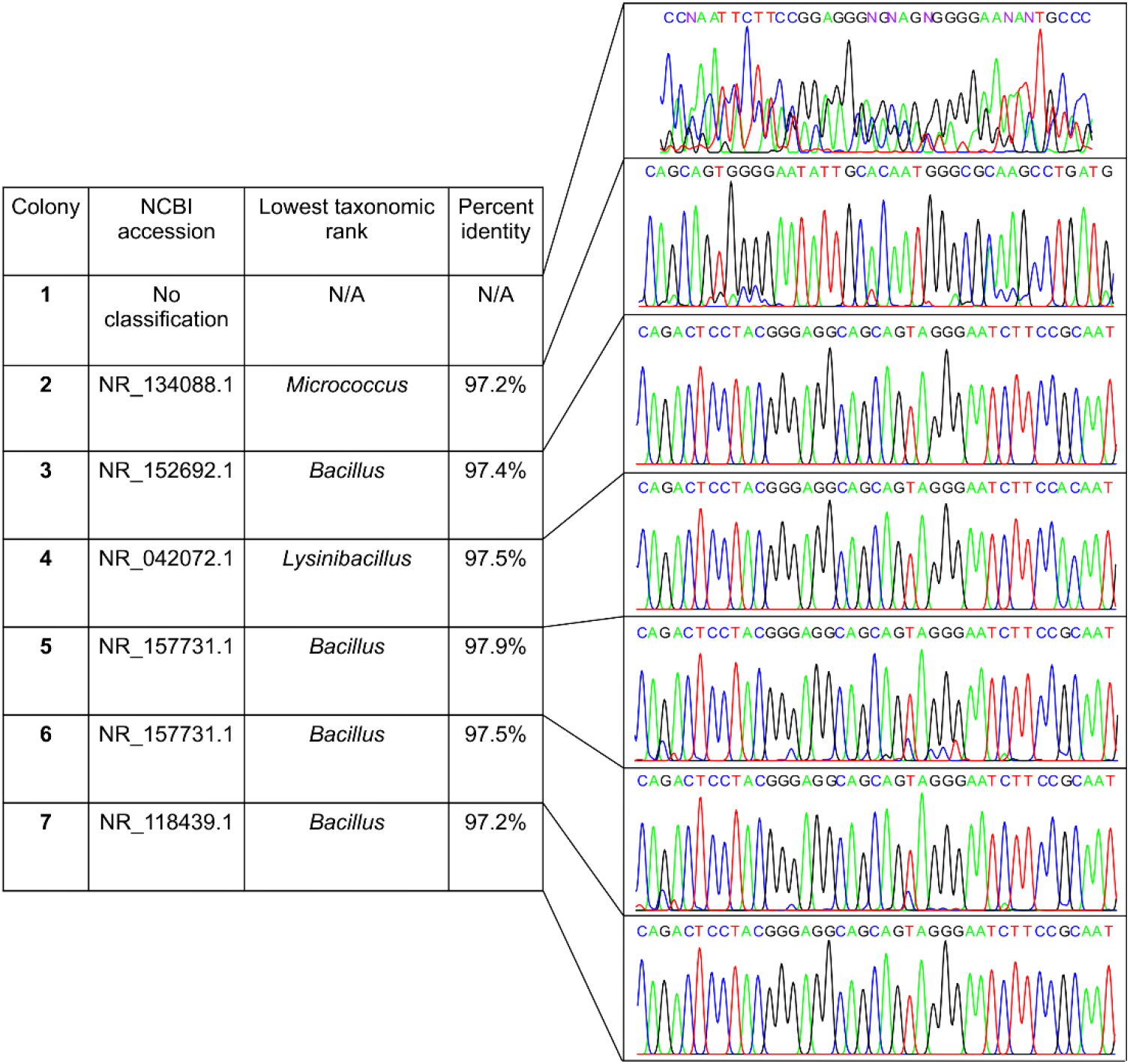
Classification of bacterial colonies by Sanger sequencing of 16S amplicons. The sequence obtained for each colony (designated **1**-**7**) was compared against NCBI database and the top match was used for taxonomic classification. For each colony, subsections of electropherograms (corresponding to base pairs 256 – 297) are shown on the right. Colony 1 was not successfully classified. The electropherograms were visualized using the R package *SangerSeqR*.

### 16S sequencing and colony identification: Nanopore

Next, we conducted MinION nanopore sequencing of full-length 16S amplicons obtained from the same seven bacterial colonies. First, we compared the performance between standard Fast basecalling model and the more recently introduced, highly discriminatory Bonito (“super accuracy”) basecalling model in converting the flow cell-generated ionic current data into sequence of nucleotide bases. Fast and Bonito basecalling of ionic current data generated a total of 133,149 and 163,552 read sequences, respectively. Table 1 summarizes the outcome for genus level classification of both basecalling methods, including the number of reads, the percentage of total reads that were successfully classified, and the mean percent identity (Ī) of all sequences for each colony. We observed that switching from Fast to Bonito basecalling while maintaining a constant Quality Score (Qscore) of 13, leads to an increase in the number of total reads and Ī, but the percentage of correct classification remains comparable. However, implementing the Bonito basecalling at a Qscore threshold of 13 and increasing the identity threshold (I) to 95%, considered to be optimal for taxonomic identification at the genus level,^25^ we noticed a substantial improvement in percentage of correct classification (≥99.4% reads were correctly classified). It is to be noted that this improvement in the classification accuracy was associated with a considerable drop in the total read number.^25^

**Table 1.** Genus-level classification of 16S amplicons for bacterial colonies **1**-**7** after nanopore sequencing. The outcomes for Fast and Bonito basecalling, and assignment of different Qscore are compared.

| Colony | Fast basecalling |  |  | Bonito basecalling |  |  | Bonito basecalling |  |  |
| --- | --- | --- | --- | --- | --- | --- | --- | --- | --- |
|  | Q ≥ 7 |  |  | Q ≥ 7 |  |  | Q ≥ 13, I ≥ 95% |  |  |
| | Total | Classified (%) | $\bar{I}$ (%) | Total | Classified (%) | $\bar{I}$ (%) | Total | Classified (%) | $\bar{I}$ (%) |
| **1** | 19,270 | 89.5 | 86.2 | 24,753 | 89.6 | 91.4 | 2,271 | 99.4 | 95.9 |
| **2** | 14,037 | 90.7 | 86.9 | 18,517 | 91.1 | 92.0 | 1,894 | 99.7 | 96.2 |
| **3** | 12,063 | 96.0 | 87.5 | 15,469 | 93.9 | 92.5 | 1,961 | 99.6 | 96.6 |
| **4** | 24,741 | 91.9 | 86.6 | 28,734 | 92.7 | 92.1 | 3,345 | 99.9 | 96.3 |
| **5** | 15,598 | 96.0 | 87.4 | 19,182 | 93.9 | 92.5 | 2,543 | 99.7 | 96.6 |
| **6** | 18,471 | 96.0 | 87.3 | 21,534 | 93.7 | 92.4 | 2,910 | 99.8 | 96.6 |
| **7** | 28,969 | 94.3 | 87.0 | 35,363 | 92.6 | 92.5 | 4,022 | 99.8 | 96.6 |

The total numbers of reads, the percentage of reads successfully classified, and the mean percent identities are presented in the table. Q thresholds of 7 and 13 correspond to an 85% and a 95% chance that each base is accurate, respectively. The classification was performed using the NCBI 16S database for reference. Abbreviations: Quality Score (Q), Percent Identity (I), Mean Percent Identity ( $\bar{I}$ )

The Bonito-basecalled sequences (Q ≥ 13, I ≥ 95%) were taxonomically identified using the EPI2ME 16S workflow, which utilizes the NCBI 16S database as a reference database (Figure 3). Calculation of relative abundances from these classified sequences showed a single bacterial genus for colonies **3**-**7** (relative abundance ≥ 99.5%) and their identities matched well with the findings from Sanger sequencing (Figure 3). In the case of colony 1, for which Sanger sequencing failed to assign a taxonomic classification with an inferior quality electropherogram, nanopore sequencing indicated the presence of two dominant taxa, namely *Alkalihalobacillus* (87.1%) and *Kocuria* (10.9%). For colony **2** also, nanopore sequencing showed the presence of two bacteria, namely *Micrococcus* (68.4%) and *Paraburkholderia* (27.7%). Interestingly, Sanger sequencing of the colony classified it as *Micrococcus* with a high 97.4% identity and had an electropherogram with clean, distinct peaks. Thus, not only the less abundant bacteria in colony **2** was not identified by Sanger sequencing but also the potential presence of a second bacterial species was not suggested by the electropherogram, or the sequence obtained.

**Figure 3.**
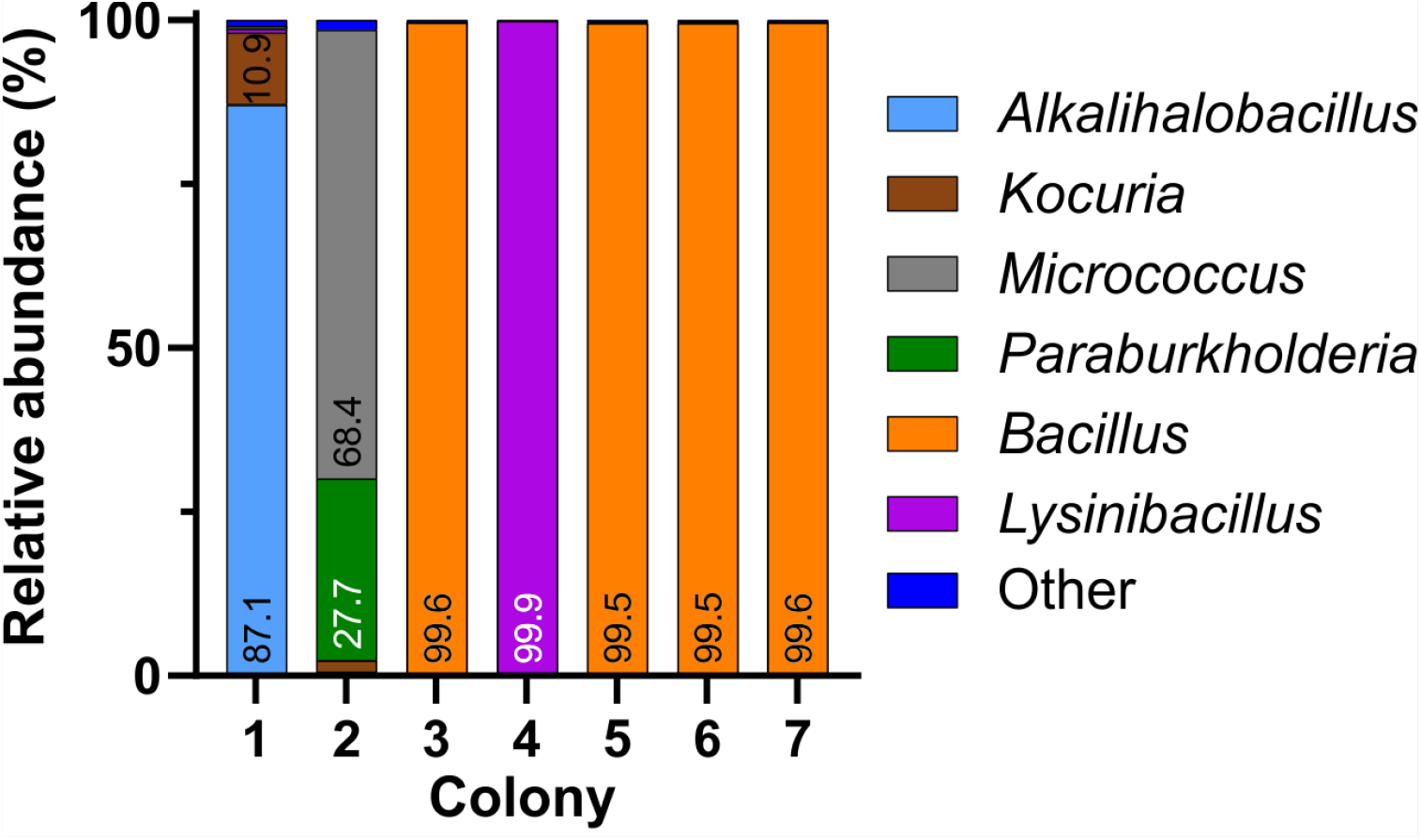
Relative abundances of bacterial genera in colonies **1**-**7**, obtained by nanopore sequencing of 16S rDNA amplicons (Bonito basecalling; Qscore ≥ 13, Percent Identity ≥ 95%).

### Evaluation on mock multispecies colonies constructed by pure bacterial DNA

Bioaerosol-derived colonies, identified through sequencing indicated the potential existence of multispecies colonies. Although Sanger sequencing of the 16S gene successfully identified typical colonies formed by a single bacterial species, we found inconsistent outcomes (percent identity and electropherogram quality) for multispecies colonies identified by nanopore sequencing.

To further understand how 16S Sanger sequences change when more than one bacterial genomic DNA is present in the sample, we designed a setup of mock multispecies colonies by mixing pure DNA samples of two bacterial species at known proportions. Pure genomic DNA of *Acinetobacter baumannii* (*A. baumannii*, ATCC 19606D-5) and *Stenotrophomonas maltophilia* (*S. maltophilia*, ATCC 13637D-5) were mixed at 3 different ratios (w/w) of 9:1, 1:1, and 1:9; Sanger sequencing was conducted in these samples and the results were compared against the pure DNA controls of *A. baumannii* and *S. maltophilia* (Fig. 4). The sequences from pure DNA controls of *A. baumannii* (sample **A**) and *S. maltophilia* (sample **E**) were classified correctly when aligned against NCBI 16S reference database, although identity matches were 95.5% and 97.4%, respectively, allowing for genus level identification. The Sanger sequence generated from the sample with 90% *A. baumannii* and 10% *S. maltophilia* (sample **B**, Fig. 4) was classified as *A. baumannii* with 96.8% identity match, while the sequence generated from the sample with 10% *A. baumannii* and 90% *S. maltophilia* (sample **D**) had closest alignment with *S. maltophilia* with 81.3% identity. For the sample with 50% of both *A. baumannii* and *S. maltophilia* (control sample **C**), the Sanger sequence was found to have closest alignment with *A. baumannii* with a low 79.3% identity. Unlike samples **A, B**, and **E**, the lower identity match for samples **C** and **D** would allow the lowest taxonomic classification to the phylum level only. The electropherograms also showed distinct changes to different levels of DNA mixing (Fig. 4). The electropherogram of pure *A. baumannii* DNA (sample **A**) had clear and separated peaks while pure *S. maltophilia* DNA (sample **E**) showed some peak overlap even though the alignment of Sanger sequence showed a high percentage identity. In mixed samples, the presence of 10% *S. maltophilia* DNA made only a minimal change to the electropherogram signal of *A. baumannii* (sample **B**), however, the presence of 10% of *A. baumannii* caused a substantial degradation of the electropherogram signal of *S. maltophilia* (sample **D**). Counterintuitively though, equal mixing of each of these two DNA (sample **C**) resulted in a relatively clean electropherogram although the identity match of the Sanger sequence obtained from the electropherogram was lower than other two mixed samples.

**Figure 4.**
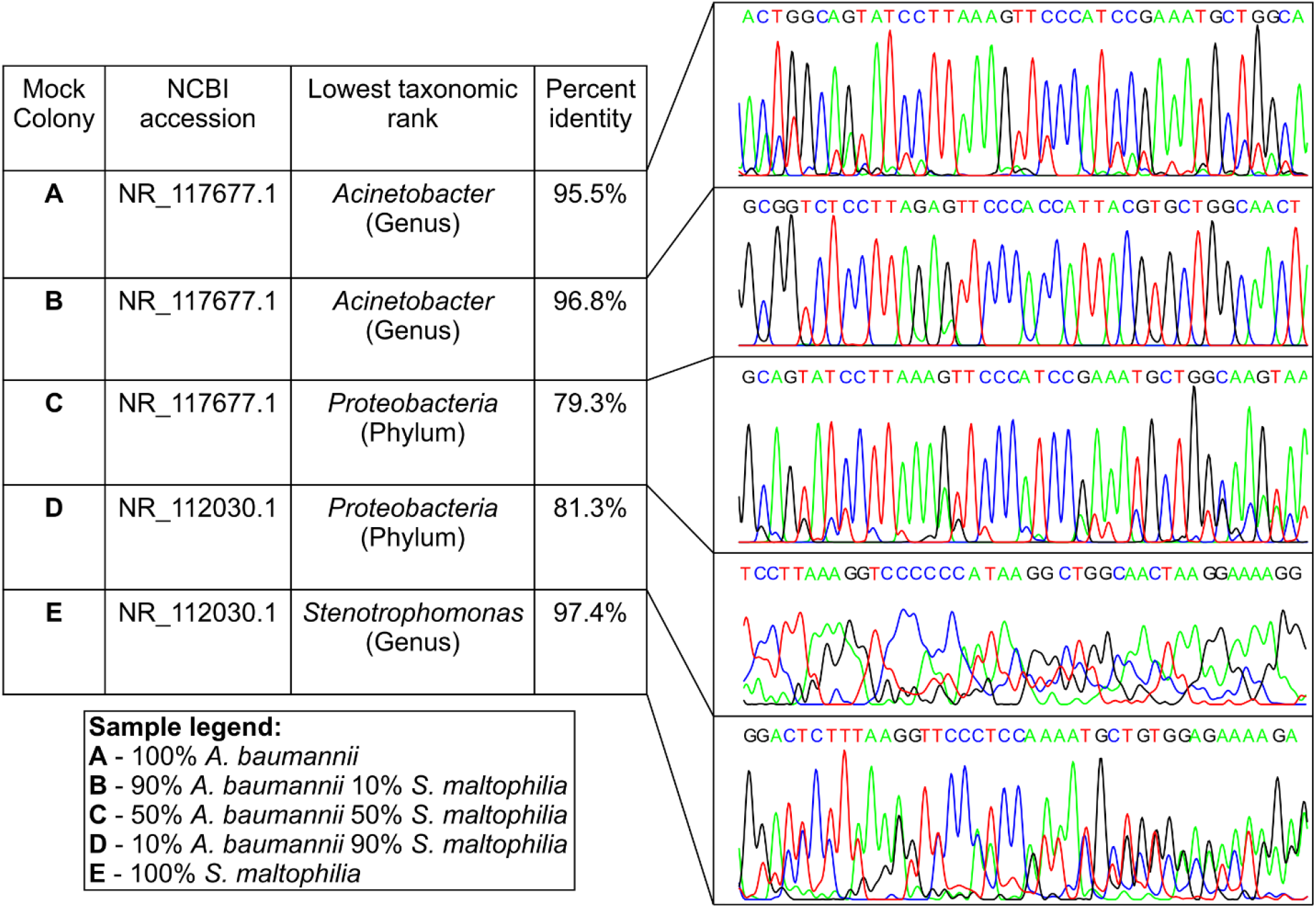
Classification results of the mock colony samples **A**-**E** by Sanger sequencing of 16S amplicons. Electropherogram segments corresponding to each sample are shown. Taxonomic classification was obtained by the best match of a sequence against the NCBI 16S database. The electropherograms were obtained using the R package *SangerSeqR*.

In parallel to Sanger sequencing, we performed nanopore sequencing of the samples from mock colonies. The Bonito-basecalled reads correctly classified >99% of reads at the genus level (Qscore ≥ 13, I ≥ 95%) and the performance was comparable for all samples irrespective of different proportions of DNA mixing. We found that the nanopore sequencing was able to successfully classify pure DNA samples (samples **A** and **E**) as well as the mixed DNA samples (samples **B, C**, and **D**) with the relative abundances reflecting the proportion of mixing (Fig. 5).

**Figure 5.**
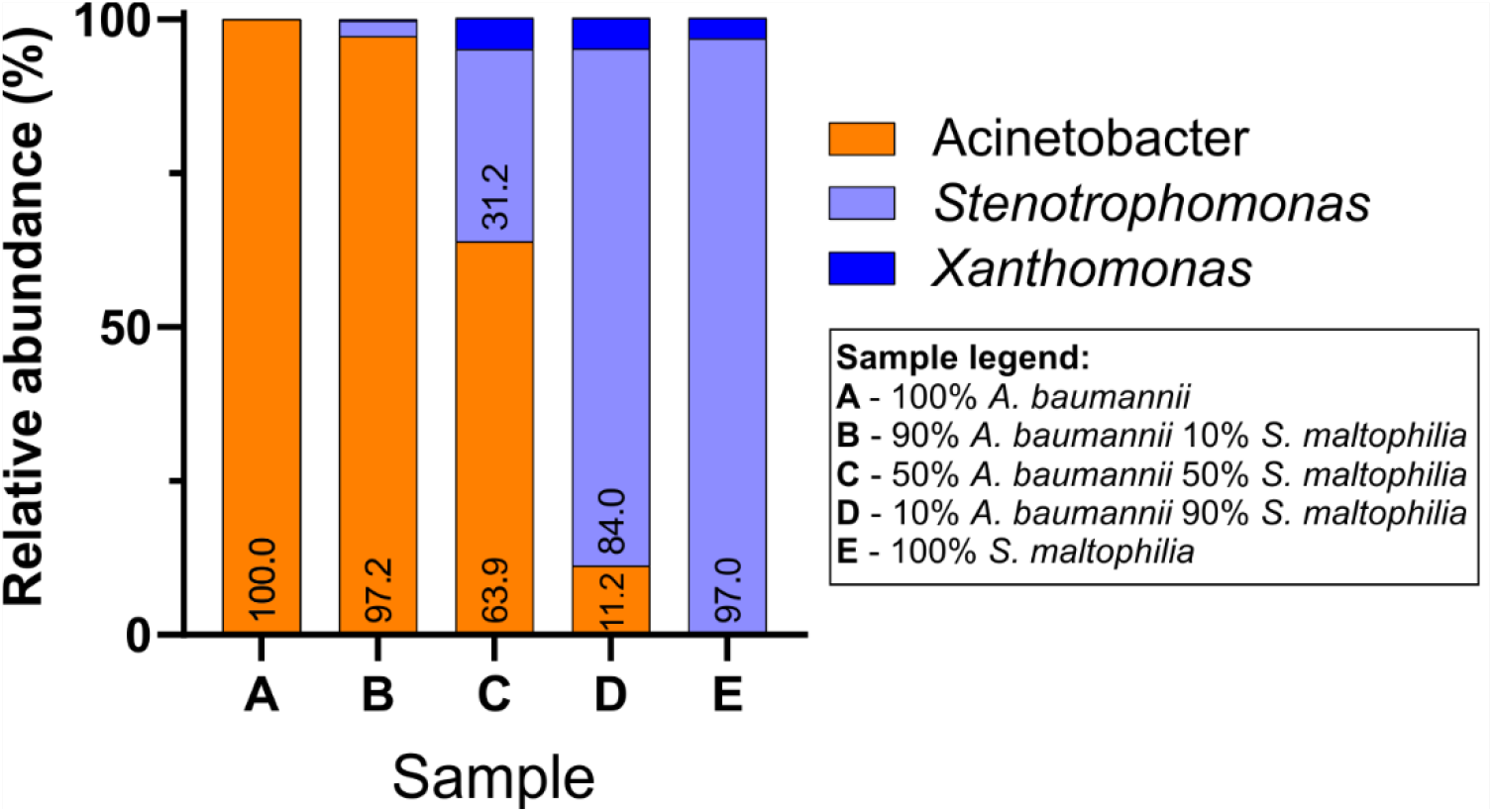
Relative abundances of bacterial genera in mock colony samples **A**-**E**, obtained by nanopore sequencing of 16S amplicons (Bonito basecalling; Qscore ≥ 13, Percent Identity ≥ 95%).

Since the species level information of pure DNA samples were known for the control experiment, we attempted species-level identification for the sequences raising the threshold of I to ≥ 99%.^35^ The new threshold criteria reduced the total number of passed reads to 5,148 (Table 2). We found that for the pure *A. baumannii* and *S. maltophilia* samples, 97.4% and 92.1% of the correctly classified sequence reads from nanopore sequencing accurately matched to the species level. A similar degree of accuracy was maintained in the species-level identification of mixed samples.

**Table 2.** Summary classification of 16S amplicons for mock colony samples **A**-**E** sequenced using nanopore sequencing (Bonito basecalling; Qscore ≥ 13). For genus and species-level classification, the thresholds used for I were set at ≥ 95% and ≥ 99%, respectively.

| Mock colony | Total | Bonito genus-level |  |  |  | Bonito species-level |  |  |  |  |
| --- | --- | --- | --- | --- | --- | --- | --- | --- | --- | --- |
| | | Q $\geq 13$, I $\geq 95\%$ | | | | Q $\geq 13$, I $\geq 99\%$ | | | | |
|  |  | CC (%) | MC (%) | UC (%) | I (%) | Total | CC (%) | MC (%) | UC (%) | I (%) |
| **A** | 41,158 | 99.9 | <0.1 | 0.1 | 97.3 | 573 | 94.1 | 2.5 | 3.4 | 99.2 |
| **B** | 79,618 | 99.7 | <0.1 | 0.3 | 97.2 | 1,419 | 86.2 | 13.6 | 2.2 | 99.3 |
| **C** | 73,554 | 99.6 | <0.1 | 0.4 | 97.3 | 1,266 | 88.2 | 10.2 | 1.6 | 99.3 |
| **D** | 84,019 | 99.9 | <0.1 | 0.1 | 97.3 | 1,233 | 91.3 | 4.0 | 4.7 | 99.2 |
| **E** | 40,366 | 99.6 | <0.1 | 0.4 | 97.3 | 657 | 85.5 | 5.0 | 9.5 | 99.2 |

The table shows the total number of reads along with the percentage of reads Correctly Classified (CC), Misclassified (MC), and Unclassified (UC) for both genus and species-level classification. The classification was performed using the NCBI 16S database for reference. Abbreviations: Qscore (Q), Percent Identity (I), Mean Percent Identity (I)

### Generation of consensus amplicon sequence from nanopore data

Although nanopore sequencing demonstrates an advantage over the Sanger method on amplicon-based identification of bacteria from multispecies colonies, one criticism of this technique is the relatively lower read accuracy of individual amplicons. For a DNA sample from a single species, this limitation is addressed through the generation of a consensus sequence from the read sequences with read accuracy being comparable to one obtained through Sanger method.^20,36^ Such approach has recently been expanded in NGSpeciesID workflow for mixed DNA samples through implementation of appropriate clustering and polishing strategies.^20^ Implementing this method on nanopore sequence data obtained from pure DNA controls of *A. baumannii* and *S. maltophilia* generated consensus sequences that accurately matched to the NCBI database sequence with > 99.9% identity (Table 3). Furthermore, the alignment to the source strain sequences (ATCC 19606D-5 and ATCC 13637D-5 for *A. baumannii* and *S. maltophilia*, respectively) matched by 99.93% for both samples, indicating a deviation of only one nucleotide the in the entire 16S region. For mock multispecies colony with 50% presence of each of these two bacterial DNA (sample **C**), two consensus sequences were generated with one having 100% match with *A. baumannii* and the other having 98.04% match with *S. maltophilia*. When applied for mock colony samples with 90% DNA from *A. baumannii* (sample **B**) or *S. maltophilia* (sample **D**), the technique returned a single consensus sequence for *A. baumannii* and *S. maltophilia*, respectively, with identity match of > 99.9%. These results confirm that highly accurate consensus sequences of 16S amplicons can be obtained from nanopore reads even in mixed colony samples, except when the relative abundance of a bacterial species is disproportionately lower than the dominant species.

**Table 3.**
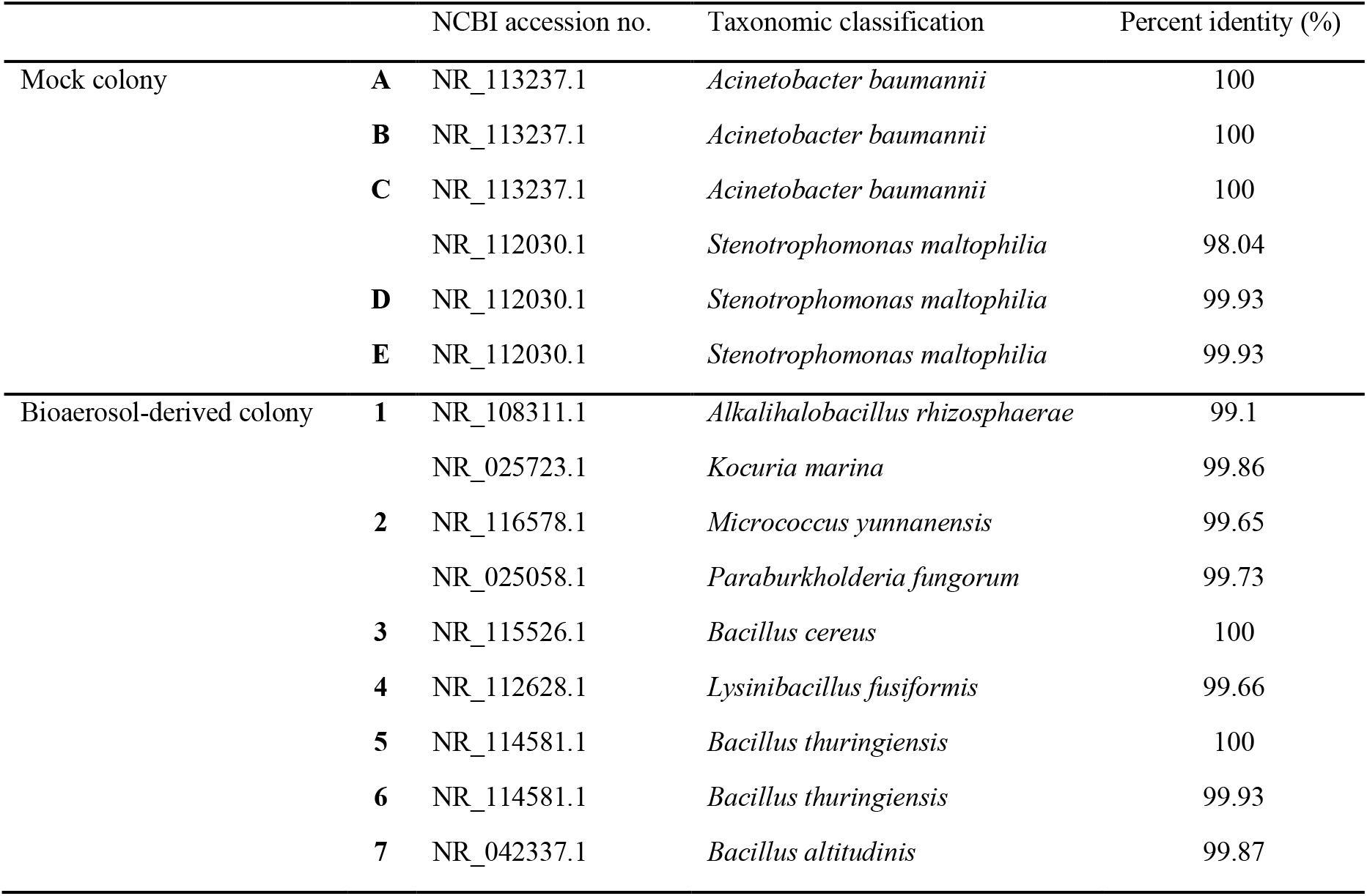
Classification of consensus sequences generated from nanopore reads of samples from mock and bioaerosol-derived colonies (Bonito basecalling; Qscore ≥ 13), obtained by alignment against NCBI 16S database using BLASTn tool.

After successful construction of consensus sequences of 16S amplicons from mock colonies, we aimed to implement it to the bioaerosol-derived colonies. The consensus sequences obtained were found to have >99% identity match with sequences at NCBI 16S database, enabling species level identification. Moreover, for colony **1** and **2**, consensus sequences of both bacteria that contributed to each of these mixed colonies, could be generated maintaining similarly high identity match to the NCBI database. It is to be noted that the total number of nanopore sequence reads for bioaerosol-derived colonies were substantially lower than the reads from the mock colonies (compare the Q ≥ 13, I ≥ 95% read numbers in Tables 1 and 2) but that didn’t adversely impact the quality of the consensus sequence. Taken together, our results indicate that16S reads from nanopore sequencing could be utilized not only to identify individual bacteria from mixed bacterial colonies and estimate their proportional abundance but also to construct a full-length amplicon sequence with high accuracy.

## Discussion

In this study, we compared Sanger and nanopore sequencing for the application of identifying single bacterial colonies using full-length 16S amplicons. We observed that both techniques are effective and corroborate well with genus-level identification when the colonies have a homogeneous composition of a single taxon. For multispecies colonies, which we encountered in two of the seven colonies of bioaerosol-derived culture, nanopore sequencing was able to identify the bacterial components forming the colony with a close estimate of their compositional representation. Furthermore, consensus sequences constructed from the nanopore reads yielded highly accurate sequence alignment (≥ 99%), enabling species level identification. Although Sanger sequencing-based identification is not suitable when multiples DNA sequences are present in a sample, we found that the sequence identity match and electropherogram quality were also less informative metrics to rule out the potential existence of multiple bacterial species in a colony.

While several studies have compared nanopore sequencing with NGS such as Illumina sequencing,^37–39^ relatively few works are directed toward the comparison of MinION sequencing and Sanger sequencing.^40^ The interest in using nanopore as a potential alternative sequencing tool to Sanger sequencing are primarily for amplicon-based assays, commonly used in forensic genetics or tracking species in the field.^36,40^ Since Sanger and nanopore sequencing are based on completely different technologies and generate a different form of output, the comparison is not straightforward and done through the generation of a consensus sequence from nanopore read sequences.^20^ Among the few comparisons being made for targeted amplicon analysis, Vasiljevic *et al*. reported using nanopore sequencing to identify animal species via the species-diagnostic region of mitochondrial cytochrome b (mtDNA cyt b) gene with an amplicon length of approximately 421 bp.^41^ Their results showed that the consensus sequences derived from nanopore sequencing were remarkably close to Sanger sequences with a deviation of not more than 1 bp. It is to be noted that the performance of Sanger sequencing starts to decrease for longer sequence lengths >1000 bases), and therefore, could have less accuracy in sequencing full-length 16S gene (∼1,500 bp).^42^ Not surprisingly, we found that Sanger method-derived 16S sequences had <98% maximum identity match against NCBI database for both pure bacterial DNA and cultured colonies, preventing a species level classification. The nanopore sequencing technology, however, is not constrained by such limitations as the read accuracy is independent of the length of DNA fragments sequenced. This is further supported by our findings that 16S consensus sequences generated from nanopore reads of pure DNA samples of *A. baumannii* and *S. maltophilia* strains are deviated by only one base from the maximally aligned sequences in the NCBI 16S database.

Sanger sequencing is still being considered a default choice in many fields for the identification and comparison of homogeneous genetic material, particularly when amplicons of sub-thousand base pairs are used for characterization.^19^ The technique can also be applied to mixed microbial samples through colony culture and isolation of individual taxa from distinct colonies. In this work, we found that Sanger sequencing of 16S amplicon successfully identified to the genus level the bioaerosol-derived colonies **3**-**7** and pure DNA control samples with an identity match of ≥ 95%. For colony **1**, the low identity match and a noisy electropherogram with overlapping peaks raised the suspicion of the presence of more than one bacterial species; subculturing followed by repeat Sanger sequencing was needed for bacterial identification of this sample. Colony **2** was identified as *Micrococcus* by Sanger sequencing with an identity match of 97.2% and was associated with a clean electropherogram with distinct peaks comparable to electropherograms from colonies **3**-**7**. However, nanopore sequencing of the sample revealed that almost one-third of the amplicons are from taxonomically distant *Micrococcus*. The mock colony experiment using pure bacterial DNA further confirmed that the presence of additional taxa in a sample has a variable impact on electropherogram quality and percent identity match for the dominant taxa, and such effects are taxa specific. This was further confirmed by the control experiments with the mix of pure DNA samples from two different bacterial species. We observed that the presence of 10% *S. maltophilia* in *A. baumannii* genomic DNA had minimum impact on the percent identity match for *A. baumannii*, however, the presence of 10% *A. baumannii* in *S. maltophilia* genomic DNA resulted in a drastic reduction in the identity match for *S. maltophilia* (97.4% to 81.3%) along with a conspicuous drop in the electropherogram quality. These results together suggest that Sanger method has a limited ability to discriminate from 16S amplicons whether a bacterial colony is a true homogeneous colony (as in colony **3**-**7**) or a multispecies colony (as in colony **2**). This uncertainty can pose a serious limitation on the applicability of Sanger sequencing on colony identification when there is potential for the existence of multispecies colony formation.^42^

Nanopore sequencing technology is emerging fast in the landscape of sequencing and is being applied for an increasing range of applications, including whole genome sequencing, microbiome, and transcriptome analysis.^31,33,34,43^ Underlying the rapid growth and increased acceptance of this technology is a continual advancement in the sequencing platform and basecalling algorithm, increased availability of protocols and bioinformatics tools for analysis, along with benchmarking studies confirming the robustness and reliability of performance.^44^ Indeed, when comparing the preexisting Fast basecalling with more recently introduced Bonito basecalling, we observed a substantial improvement in the quality of sequence reads (Īincreased by ∼5%) along with a larger number of reads passing a preset quality threshold. Implementing appropriate quality threshold (Qscore ≥ 13, I ≥ 95%), we were able to achieve accurate genus-level classification. However, the biggest advantage of nanopore sequencing over the Sanger method in the context of colony identification comes from its ability to identify all bacterial taxa present in the colony as in the case of colonies 1 and 2. Additionally, the control experiment with pure genomic DNA demonstrated that the relative abundance of bacteria observed by the nanopore sequencing reasonably reflects their proportion in a mixed sample, although some deviation can result from the variability in 16S gene copies present among bacterial species.^14,45^

The capability of long-read by MinION nanopore or PacBio sequencing platform offers a clear advantage over short-read sequencing technologies for applications in taxonomical identification or classification through targeted gene amplification. For the 16S gene, sequences longer than 1,300 bases are considered to be suitable for reliable results.^15^ However 16S taxonomical classification by Illumina-based sequencing usually targets the hypervariable regions V4, V3-V4, or V4-V5 of the 16S gene due to the limitation of this technique to read only a short span of the 16S sequence.^26,27^ Such restriction imposed on amplicon length allows the identification only up to the genus level. While near full-length 16S sequencing on the Illumina platform has been achieved by using unique, random sequences to tag individual 16S gene templates, the long, complex procedure is not practical for routine implementation.^46^ The long-read sequencing enables the analysis of full-length 16S gene amplicon and such coverage has been shown to successfully identify microbiota to species level resolution.^27^ A recent study using PacBio long-read sequencing platform could even achieve the read accuracy to single nucleotide level through the construction of circular consensus sequences (CCS) followed by implementation of advanced algorithm for analysis that enabled strain level identification.^47^ Although this result is highly accurate, the need for an expensive equipment, and a relatively complex sample preparation and analysis process along with a higher cost associated for sequencing make it less suitable for routine identification of bacterial colonies. In our work, we found that implementation of Bonito basecalling followed by selection of high-quality reads (Qscore ≥ 13, I ≥ 99%) enabled successful species-level classification of 97.4% and 92.1% of the amplicon sequences derived from pure DNA samples of *A. baumannii* and *S. maltophilia*, respectively. It is to be noted that ∼99% of identity threshold is recommended for species level identification from full-length 16S sequence, and such high threshold leads to the rejection of a significant proportion of reads, and therefore, requires a higher number of raw reads for analysis. Our results show that construction of consensus sequence would be an attractive alternative strategy for species level identification where highly accurate (>99%) identity match was observed even for mixed colony samples.^35^ Moreover, the quality of the of the consensus sequence was maintained even with a smaller number of reads per sample (as observed with bioaerosol-derived colonies), which could be highly advantageous in reducing the sequencing cost by enabling a larger number of samples to be sequenced per flow cell or utilization of flongle, a more affordable option for nanopore sequencing. One limitation in the implementation of this technique would be potentially missing a bacterial species in a multispecies culture when its relative abundance is very low in comparison to the dominant species.

Microbes in the air can exist both as a single organism and as aggregates, often bound to particulate matter.^23,24^ Due to their low numbers in air, bioaerosol samples are usually plated directly on agar media for culture-based assays to study viable microorganisms. The aggregates of viable bacteria can potentially lead to the formation of multispecies colonies, and therefore, such possibilities should be considered when analyzing single colonies. Our results show the advantage of nanopore sequencing over Sanger sequencing for accurate identification of bacterial taxa in such samples. With the improvement of sequencing accuracy in recent years, the ability to conduct sequencing in the lab with the relatively inexpensive MinION device, the availability of a streamlined protocol for 16S analysis, and the possibility to multiplex multiple samples, nanopore sequencing could be a powerful sequencing tool for culture-based assays of bioaerosol or other complex environmental samples.

## Methods

### Bioaerosol Sample Collection and Bacterial Culture

Bioaerosol samples were collected from the Clarkson University campus using a portable bioaerosol sampler (Trace Aerosol sensor and Collector for Bio-particles, TracB), which captured airborne particles on a collection plate by electrostatic precipitation.^48^ After two weeks of sample collection by the TracB device, particles deposited on the collection plate were wiped using a sterile cotton swab, pre-wetted with sterile PBS. The sample was immediately inoculated on Tryptic Soy Agar (TSA, Difco) plates by gently spreading the swab over the agar media. Following sample inoculation, TSA plates were incubated at 30 °C for 16-18 h to promote bacterial growth and colony formation before further analysis.

### DNA Extraction from Bacterial Colonies

Bacterial colonies formed on TSA plates following inoculation with bioaerosol samples were used for genomic DNA extraction. Bacteria from a single, distinct colony was carefully picked up by a sterile inoculation loop, and DNA extraction was performed using the FastDNA SPIN Kit for Soil (MP Biomedicals #116560200) following the manufacturer’s protocol. The extraction process included a bead beating step using MP Biomedicals FastPrep-24™ bead beater, operating at 6.0 m/s for 1 min. The DNA was eluted in 50 µL volume. The quality (260/280 ratio) and concentration of extracted DNA were measured using a Nanodrop spectrophotometer (ThermoFisher Scientific) and a Quantus Fluorometer with Quantifluor ONE dsDNA dye (Promega #E4891).

### Preparation of Mock Colony Samples

The mock colony samples were prepared using pure genomic DNA from two known control bacterial taxa, namely *Acinetobacter baumannii* strain 2208 (ATCC 19606D-5) and *Stenotrophomonas maltophilia* strain 810-2 (ATCC 13637D-5). *A. baumannii* and *S. maltophilia* DNA (0.5 ng/µL in nuclease free water) were mixed at predetermined volumetric ratios (9:1, 1:1, and 1:9) to prepare mock multispecies colony samples with different proportional abundance of these two DNA. For example, a sample with 90% *A. baumannii* and 10% *S. maltophilia* was prepared by mixing 4.5 µL of *A. baumannii* DNA with 0.5 µL of *S. maltophilia* DNA. The mixed DNA or pure bacterial DNA were further used for Sanger and nanopore sequencing as mock colony samples.

### Sanger Sequencing

For Sanger sequencing, full-length 16S rRNA gene (spanning hypervariable regions V1 to V9) from genomic DNA samples of bioaerosol-derived and mock colonies samples was amplified through PCR using 27F (5’-GAGTTTGATCATGGCTCAG-3’and 1492R (5’- ACGGCTACCTTGTTACGACTT-3’) primers, purchased from Invitrogen. The PCR reaction mixture contained: 12.5 µL LUNA LongAmp Taq 2x Master Mix (NEB M0287S), 5.5 µL Nuclease-Free Water (ThermoFisher Scientific #AM9937), 5 µL of bacterial DNA, and 1 µL of the 27F and 1492R primers per reaction. The PCR cycling conditions used were as follows: initial denaturation at 95 °C for 1 min, followed by 26 cycles of denaturation at 95 °C for 20 s, annealing at 55 °C for 30 s, and extension at 65 °C for 2 min; this was followed by a single 5 min extension at 65 °C. After the completion of PCR, the amplification products were run through 1.2% agarose gel electrophoresis to confirm the presence of a single band at ∼1500 bp. 400-600 ng of the PCR amplification products were sent to Eurofins Genomics for Sanger sequencing using the tube sequencing format. The Sanger sequences returned by Eurofins Genomics in a fasta file format were aligned against NCBI 16S reference database for classification using BLASTn algorithm. The electropherogram files were visualized using the package *SangerSeqR (https://bioconductor.org/packages/release/bioc/html/sangerseqR.html)* in R software.

### Nanopore Sequencing

16S rRNA gene amplification and barcoding for nanopore sequencing were performed using the 16S Barcoding Kit 1-24 (SQK-16S024) from Oxford Nanopore Technologies (ONT) following a protocol recommended by the manufacturer. The kit contained barcoded full-length 16S rRNA gene primers (9/27F and 1492R) to be used for PCR amplification. The PCR reaction mixture contained 25 µL LUNA LongAmp Taq 2x Master Mix CAT#M0287S, 5 µL Nuclease-Free Water (ThermoFisher Scientific #AM9937), 10 µL bacterial DNA, and 10 µL of barcoded16S primers. The following PCR cycling conditions were used: initial denaturation at 95 °C for 1 min; 26 cycles of denaturation at 95 °C for 20 sec, annealing at 55 °C for 30 sec, and extension at 65 °C for 2 min; a single 5 min extension at 65 °C. The PCR product (barcoded 16S amplicon) was cleaned up from the PCR reaction mixtures using AMPure XP Solid Phase Reversible Immobilization (SPRI) paramagnetic beads (Beckman-Coulter #A63880). 50 µL of the PCR reaction mixture containing the amplified product was mixed with equal volume of SPRI bead suspension and a magnetic separation rack was used to separate DNA-bound beads from rest of the solution. After two washes with 70% ethanol, the cleaned PCR product was eluted from the beads in 10 µL of buffer solution containing 10 mM Tris-HCl pH 8.0 with 50 mM NaCl and the DNA concentration was quantified on a Quantus Fluorometer using Quantifluor ONE dsDNA dye (Promega #E4891).

The nanopore sequencing of barcoded 16S amplicons were performed either using a Flongle™ or a MinION™ flow cell (R.9.4.1). For the sequencing run on Flongle, the flow cell was first primed with a mix of 117 µL of flush buffer and 3 µL of flush tether to wash out the storage buffer solution. Once flushed, the flow cell was loaded with a solution containing 5 µL of DNA amplicon library (premixed with 0.5 µL rapid adapter protein), 15 µL of sequencing buffer, and 10 µL of library loading beads, after which the sequencing run was started. To conduct sequencing on a MinION flow cell, it was first primed by loading 800 µL of flush buffer / flush tether mix through the priming port and incubating for 5 min. Immediately before loading the DNA library another 200 µL of flush buffer / flush tether mix was added to the priming port with the Spot-On port open. Sequencing was started after adding a solution mix containing 11 µL of sample DNA library (previously mixed with 1 µL of the rapid adapter protein), 34 µL of sequencing buffer, 4.5 µL of nuclease-free water, and 25.5 µL of the loading beads (added immediately before use), and through the Spot-On port.

### Basecalling & Sequence Identification

Basecalling was performed using Guppy Basecalling Software (version 5.0.7+2332e8d65) from ONT.^49^ The more recently released Bonito “super accuracy” basecaller model was used alongside the Fast basecaller model to compare the basecalling performance on the accuracy of nanopore read sequences.

Nanopore read sequences were identified using the EPI2ME 16S analysis pipeline (EPI2ME Fastq 16S v21.03.05), which performs Nucleotide Basic Local Alignment Search Tool (BLASTn) on each individual reads against the NCBI 16S reference database. For taxonomical classification, minimum identity threshold for species and genus level were set at 99% and 95%, respectively.^25^ Additionally, only reads that returned lowest common ancestor (lca) value of 0 during EPI2ME alignment were considered successfully classified and sequences with lca value of -1 or 1 were considered unclassified. For mock colony samples, the classified sequences were further separated into correctly classified and misclassified category based on whether the classification correctly matches with the known references.^39^ The classification of Sanger sequences was also performed by alignment the NCBI 16S reference database using the BLASTn tool. The minimum identity threshold criteria used for taxonomic classification were same as used for nanopore sequences. Additionally, the top three alignment matches were examined for agreement to assign to a specific taxon (e.g., the top three alignment matches should be from same genus for genus-level assignment). If an agreement was not reached, then the assignment was provided to lowest common ancestor.

### Construction of Consensus Sequence from Nanopore Reads

Consensus sequences from nanopore reads were constructed using NGSpeciesID program.^20^ NGSpeciesID utilized .fastq files from guppy basecalling to The program creates one or a few highly accurate representative sequences from thousands of nanopore reads, by using a combination of selective clustering and polishing strategies. Bonito basecalled .fastq files of 16S reads were used as input and the intended target length parameter was set as 1500 (approximate length of 16S gene) along with the maximum deviation from target parameter was set as 500 to cluster and generate consensus sequences of 16S amplicons for all bacterial species present in the sample.

## Conclusions

DNA sequencing of PCR-amplified targeted gene to detect microbial colonies is routinely practiced in microbiological laboratories and commonly utilizes the Sanger method. In this study, we explored the potential of nanopore sequencing for the purpose of comparing it against the Sanger technique. Evaluating full-length 16S gene-based identification of bioaerosol-derived bacterial colonies and mock colonies, we found that Sanger sequencing provides consistent genus level classification for single-species colonies; for multispecies colonies, the sequencing method is not only ineffective for identification but also cannot reliably indicate such possibility. Nanopore sequencing of the same set of samples successfully identified both single- and multispecies colonies along with providing approximate relative abundance of the bacterial taxa in the multispecies colonies. Furthermore, species level identification was shown to be possible by construction of highly accurate consensus sequences from a small number of sequence reads. Thus, our findings suggest that nanopore sequencing could be an attractive alternative to accurate species level colony identification, especially for complex environmental samples such as bioaerosols, which can lead to multispecies colony formation.

## Acknowledgments

This study was supported by the funding from National Science Foundation (NSF STTR Phase II, Award No 1853522) and New York State Energy Research and Development Authority (NYSERDA). Daniel T. Fuller received support from the Lawrence ‘57 and Antoinette Delaney Ignite Research Fellowship.

## Data Availability

The datasets used in this article will be available upon request to the authors.

## References

1. Urbano, R., Palenik, B., Gaston, C. J. & Prather, K. A. Detection and phylogenetic analysis of coastal bioaerosols using culture dependent and independent techniques. Biogeosciences 8, 301–309 (2011).

2. Tiwari, A., Oliver, D. M., Bivins, A., Sherchan, S. P. & Pitkänen, T. Bathing Water Quality Monitoring Practices in Europe and the United States. Int J Environ Res Pu 18, 5513 (2021).

3. Banerjee, G. et al. Application of advanced genomic tools in food safety rapid diagnostics: challenges and opportunities. Curr Opin Food Sci 47, 100886 (2022).

4. Peker, N., Couto, N., Sinha, B. & Rossen, J. W. Diagnosis of bloodstream infections from positive blood cultures and directly from blood samples: recent developments in molecular approaches. Clin Microbiol Infec 24, 944–955 (2018).

5. Gandolfi, I., Bertolini, V., Ambrosini, R., Bestetti, G. & Franzetti, A. Unravelling the bacterial diversity in the atmosphere. Appl Microbiol Biot 97, 4727–4736 (2013).

6. Velusamy, V., Arshak, K., Korostynska, O., Oliwa, K. & Adley, C. An overview of foodborne pathogen detection: In the perspective of biosensors. Biotechnol Adv 28, 232–254 (2010).

7. Dowd, S. E. et al. Survey of bacterial diversity in chronic wounds using Pyrosequencing, DGGE, and full ribosome shotgun sequencing. Bmc Microbiol 8, 43–43 (2008).

8. Marsh, R. L. et al. Prevalence and subtyping of biofilms present in bronchoalveolar lavage from children with protracted bacterial bronchitis or non-cystic fibrosis bronchiectasis: a cross-sectional study. Lancet Microbe (2022) doi:10.1016/s2666-5247(21)00300-1.

9. Blagodatskaya, E. & Kuzyakov, Y. Active microorganisms in soil: Critical review of estimation criteria and approaches. Soil Biology Biochem 67, 192–211 (2013).

10. Fröhlich-Nowoisky, J. et al. Bioaerosols in the Earth system: Climate, health, and ecosystem interactions. Atmos Res 182, 346–376 (2016).

11. Ferguson, R. M. W. et al. Bioaerosol biomonitoring: Sampling optimization for molecular microbial ecology. Mol Ecol Resour 19, 672–690 (2019).

12. Woese, C. R. & Fox, G. E. Phylogenetic structure of the prokaryotic domain: The primary kingdoms. Proc National Acad Sci 74, 5088–5090 (1977).

13. Janda, J. M. & Abbott, S. L. 16S rRNA Gene Sequencing for Bacterial Identification in the Diagnostic Laboratory: Pluses, Perils, and Pitfalls▿. J Clin Microbiol 45, 2761–2764 (2007).

14. Větrovský, T. & Baldrian, P. The Variability of the 16S rRNA Gene in Bacterial Genomes and Its Consequences for Bacterial Community Analyses. Plos One 8, e57923 (2013).

15. Yarza, P. et al. Uniting the classification of cultured and uncultured bacteria and archaea using 16S rRNA gene sequences. Nat Rev Microbiol 12, 635–645 (2014).

16. Johnson, J. S. et al. Evaluation of 16S rRNA gene sequencing for species and strain-level microbiome analysis. Nat Commun 10, 5029 (2019).

17. Karger, B. L. & Guttman, A. DNA sequencing by CE. Electrophoresis 30, S196–S202 (2009).

18. Baudhuin, L. M. et al. Confirming Variants in Next-Generation Sequencing Panel Testing by Sanger Sequencing. J Mol Diagnostics 17, 456–461 (2015).

19. Hebert, P. D. N. et al. A Sequel to Sanger: amplicon sequencing that scales. Bmc Genomics 19, 219 (2018).

20. Sahlin, K., Lim, M. C. W. & Prost, S. NGSpeciesID: DNA barcode and amplicon consensus generation from long-read sequencing data. Ecol Evol 11, 1392–1398 (2021).

21. McCully, L. M., Bitzer, A. S., Seaton, S. C., Smith, L. M. & Silby, M. W. Interspecies Social Spreading: Interaction between Two Sessile Soil Bacteria Leads to Emergence of Surface Motility. Msphere 4, e00696–18 (2019).

22. Xiong, L. et al. Flower-like patterns in multi-species bacterial colonies. Elife 9, e48885 (2020).

23. Tham, K. W. & Zuraimi, M. S. Size relationship between airborne viable bacteria and particles in a controlled indoor environment study. Indoor Air 15, 48–57 (2005).

24. Eduard, W. Measurement methods and strategies for non-infectious microbial components in bioaerosols at the workplace. Analyst 121, 1197–1201 (1996).

25. Edgar, R. C. Accuracy of taxonomy prediction for 16S rRNA and fungal ITS sequences. Peerj 6, e4652 (2018).

26. Nelson, M. C., Morrison, H. G., Benjamino, J., Grim, S. L. & Graf, J. Analysis, Optimization and Verification of Illumina-Generated 16S rRNA Gene Amplicon Surveys. Plos One 9, e94249 (2014).

27. Matsuo, Y. et al. Full-length 16S rRNA gene amplicon analysis of human gut microbiota using MinION™ nanopore sequencing confers species-level resolution. Bmc Microbiol 21, 35 (2021).

28. Shendure, J. et al. DNA sequencing at 40: past, present and future. Nature 550, 345–353 (2017).

29. Jain, M., Olsen, H. E., Paten, B. & Akeson, M. The Oxford Nanopore MinION: delivery of nanopore sequencing to the genomics community. Genome Biol 17, 239 (2016).

30. Zascavage, R. R., Thorson, K. & Planz, J. V. Nanopore sequencing: An enrichment-free alternative to mitochondrial DNA sequencing. Electrophoresis 40, 272–280 (2019).

31. Mantere, T., Kersten, S. & Hoischen, A. Long-Read Sequencing Emerging in Medical Genetics. Frontiers Genetics 10, 426 (2019).

32. Leggett, R. M. et al. Rapid MinION profiling of preterm microbiota and antimicrobial-resistant pathogens. Nat Microbiol 5, 430–442 (2020).

33. Ciuffreda, L., Rodríguez-Pérez, H. & Flores, C. Nanopore sequencing and its application to the study of microbial communities. Comput Struct Biotechnology J 19, 1497–1511 (2021).

34. Workman, R. E. et al. Nanopore native RNA sequencing of a human poly(A) transcriptome. Nat Methods 16, 1297–1305 (2019).

35. Edgar, R. C. Updating the 97% identity threshold for 16S ribosomal RNA OTUs. Bioinformatics 34, 2371–2375 (2018).

36. Maestri, S. et al. A Rapid and Accurate MinION-Based Workflow for Tracking Species Biodiversity in the Field. Genes-basel 10, 468 (2019).

37. Heikema, A. P. et al. Comparison of Illumina versus Nanopore 16S rRNA Gene Sequencing of the Human Nasal Microbiota. Genes-basel 11, 1105 (2020).

38. Karst, S. M. et al. Retrieval of a million high-quality, full-length microbial 16S and 18S rRNA gene sequences without primer bias. Nat Biotechnol 36, 190–195 (2018).

39. Winand, R. et al. Targeting the 16S rRNA Gene for Bacterial Identification in Complex Mixed Samples: Comparative Evaluation of Second (Illumina) and Third (Oxford Nanopore Technologies) Generation Sequencing Technologies. Int J Mol Sci 21, 298 (2019).

40. Ogden, R., Vasiljevic, N. & Prost, S. Nanopore sequencing in non-human forensic genetics. Emerg Top Life Sci 5, 465–473 (2021).

41. Vasiljevic, N. et al. Developmental validation of Oxford Nanopore Technology MinION sequence data and the NGSpeciesID bioinformatic pipeline for forensic genetic species identification. Forensic Sci Int Genetics 53, 102493 (2021).

42. Krishnakumar, R. et al. Systematic and stochastic influences on the performance of the MinION nanopore sequencer across a range of nucleotide bias. Sci Rep-uk 8, 3159 (2018).

43. Jain, M. et al. Nanopore sequencing and assembly of a human genome with ultra-long reads. Nat Biotechnol 36, 338–345 (2018).

44. Leidenfrost, R. M., Pöther, D.-C., Jäckel, U. & Wünschiers, R. Benchmarking the MinION: Evaluating long reads for microbial profiling. Sci Rep-uk 10, 5125 (2020).

45. Kembel, S. W., Wu, M., Eisen, J. A. & Green, J. L. Incorporating 16S Gene Copy Number Information Improves Estimates of Microbial Diversity and Abundance. Plos Comput Biol 8, e1002743 (2012).

46. Burke, C. M. & Darling, A. E. A method for high precision sequencing of near full-length 16S rRNA genes on an Illumina MiSeq. Peerj 4, e2492 (2016).

47. Callahan, B. J. et al. High-throughput amplicon sequencing of the full-length 16S rRNA gene with single-nucleotide resolution. Nucleic Acids Res 47, e103–e103 (2019).

48. Priyamvada, H. et al. Design and evaluation of a new electrostatic precipitation-based portable low-cost sampler for bioaerosol monitoring. Aerosol Sci Tech 1–13 (2020) doi:10.1080/02786826.2020.1812503.

49. Wick, R. R., Judd, L. M. & Holt, K. E. Performance of neural network basecalling tools for Oxford Nanopore sequencing. Genome Biol 20, 129 (2019).

